# A 3D Reconstruction Algorithm for Real-time Simultaneous Multi-Source EIT Imaging for Lung Function Monitoring

**DOI:** 10.1101/2020.05.29.124222

**Authors:** Tzu-Jen Kao, Bruce Amm, David Isaacson, Jonathan Newell, Gary Saulnier, Jennifer L. Mueller

## Abstract

Monitoring regional pulmonary ventilation and pulsatile perfusion changes in a 3D region of interest (ROI) of the lung is a promising application for electrical impedance tomography (EIT). This paper describes a 3D analytical reconstruction algorithm that was embedded in a prototype EIT system to enable a real-time image reconstruction at nearly 20 frames per second for monitoring impedance changes in the chest in real-time. The derivation and results of the 3D analytical forward solution and inverse solution and details of the real-time reconstruction algorithm are given. The algorithm and EIT system are validated with simulated data, in-vitro phantoms, and finally shown to be capable of imaging ventilation and pulsatile perfusion in human subjects. The human subject data was obtained using a high-precision, high-speed and simultaneous multiple current source (SMS-EIT) developed by GE Research. Data was collected using four rows of 8 electrodes for a healthy adult male subject and 2 rows of 16 electrodes for six healthy human female subjects, with one row placed above the breasts and a second row placed at the infra-mammary fold. Each of the female subjects performed a breathing maneuver with a volumetric incentive spirometer, and the volume of air inhaled was calculated from the EIT images. Pulsatile perfusion images were computed from this data, and regional lung filling was also analyzed.

## 1 Introduction

Electrical Impedance Tomography (EIT) is an emerging medical imaging technology that estimates the internal electrical properties of the region of interest (ROI) based upon voltage measurements arising from the application of low-amplitude, low-frequency AC current applied to electrodes that are placed on the surface [Barber 1984, Newell 1988, Brown 2003, Holder 2005]. The most developed and widespread use of EIT is for pulmonary imaging, for which the literature is extensive, and we refer to the review articles [Frerichs 2017, Martins 2019, Nguyen 2012, Frerichs 2000] for more information. EIT is especially suitable for bedside pulmonary monitoring or for patients with chronic respiratory disease because it is noninvasive, non-ionizing, and an image can be produced in real time and obtained simultaneously with pulmonary function tests [Muller 2018].

The EIT reconstruction problem is to solve for the electrical conductivity and permittivity distributions inside the body based on the measured voltages and the known excitation currents, in 2D or 3D. It has long been recognized that 3D reconstruction algorithms increase accuracy and decrease artefacts caused by out-of-plane objects [Blue 2000, Eyuboglu 1989, Guardo 1991, Rabbani 1991] and are necessary to image the full lung and to obtain accurate lung volume estimates. In this work, we report the derivation of a real-time 3D reconstruction algorithm based on the ToDLeR (Three-Dimensional Linearized Reconstruction) algorithm [Blue 1997, Blue 2000], which was a simplification and a faster version of the inversion algorithm of Goble [Goble 1990]. The algorithm is implemented in the GENESIS SMS-EIT system, and the reconstructed conductivity images are produced at 20 frames per second on a 32-electrode configuration. Preliminary studies of healthy human subjects with this system have been reported [Amm 2014, Kao 2014]. In this paper we present the details of the solutions and the gradient computations that facilitate the real-time implementation. We further demonstrate the capabilities of the system and algorithm for estimating lung volumes on female subjects. A multiple electrode row 3-D reconstruction algorithm for lung imaging must be effective on both male and female subjects, and this small study demonstrates that lung volumes can be estimated from the images, regional lung filling can be seen, and images of pulsatile pulmonary perfusion can be computed from the data. This is not intended to be a comprehensive validation study.

Lung volume estimates and regional ventilation maps are important clinical measures that can be obtained from electrical impedance tomography images. Clinical validation against CT images has shown that EIT is effective for obtaining images of regional ventilation distribution [Frerichs 2002a, Frerichs 2002, Costa 2009, Smit 2004, Victorino 2004]. However, most EIT systems use a single row of electrodes placed around the chest, and very few studies have been performed on female subjects. In this work, data was collected on six healthy human female subjects with one row of 16 electrodes placed above the breasts and a second row of 16 electrodes placed at the infra-mammary fold. A breathing maneuver was performed with a volumetric incentive spirometer, and 3-D images were computed using the real-time 3D algorithm described in Section II.B. The volume of air inspired was then calculated from the 3-D EIT images using the method derived in [Muller 2015]. The results are compared to the inspiration volumes recorded by the volumetric incentive spirometer during the maneuver. Regional lung filling is also analyzed.

The paper is organized as follows. Section II.A. describes the forward problem and the analytic solution to the homogenous case. In Section II.B. we report the detailed derivation of the solution to the inverse problem based on the ToDLeR algorithm for real-time implementation. The validation on simulated data is in Section III.A. and on phantom data in Section III.B. Results on human subjects are in Section IV, with the study of lung volume estimation on the female subjects in Section IV.B.

## 2 Methods

The reconstruction algorithm is based on a perturbation approximation. It assumes that the conductivity change distribution differs only slightly from a uniform conductivity distribution. To reconstruct the conductivity, a forward solution and the Jacobian of the forward model with respect to a small perturbation in conductivity will be needed.

### 1.1 The Forward Model for EIT

The electromagnetic field introduced by the applied current on the surface is governed by Maxwell’s equations. At low current and frequency, it can be simplified to the generalized Laplace equation:

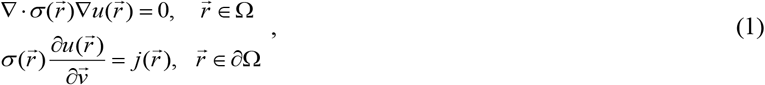

where Ω is a bounded region with boundary ∂Ω, 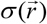 is the conductivity distribution, *u* is the electrical potential, 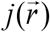 is the externally applied current density, and 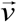 is the outward-pointing normal vector to the boundary. The forward problem in EIT is to solve for the resulting voltages on the surface of a body from the given current applied on the surface and the internal conductivity distribution.

In the homogeneous case, the first equation in (1) can be reduced to a second order partial differential equation (PDE)

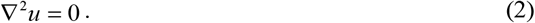

A variety of methods can be used to solve the PDE in (2), such as finite element method (FEM), which can be used to solve for complex geometries. In our targeted application of lung function monitoring, we simplified the geometry of a human torso as a cylinder, with multiple rings of electrodes placed on the side wall. With this simplification, (2) can be solved analytically using separation of variables. We include the derivation here since explicit form of the Fourier coefficients is relevant to our method. See also [Pidcock 1995, Evans 2010] for analytic solutions in this geometry.

In a cylindrical coordinate system, with subscript notation, the boundary condition in (1) can be rewritten as

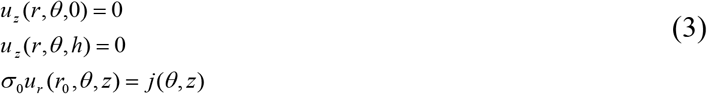

where *h* is the height of the cylinder and *r_0_* is the radius of the cylinder, and *u_z_* = ∂*u*/∂*z*. The solution to (2) with the first two boundary conditions in (3), is then

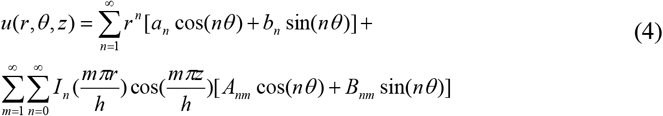

where *I_n_* is the modified Bessel function of order *n*:

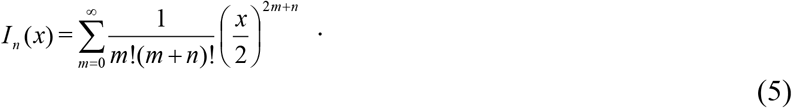

The Fourier coefficients *a_n_*, *b_n_*, *A_nm_*, and *B_nm_* are to be determined by using the third boundary condition in (3). In this study, we applied the ‘ave-gap model’ as our electrode model. In the ‘ave-gap model’, we assume that the current density is evenly distributed over the surface of each electrode and that the potential of each electrode is the average value of the potential over the area covered by the electrode. This electrode model does not take into account the shunting effect and the contact impedance between the electrodes and object [Cheng 1989]. By using this electrode model, the current density on the electrode can be approximated as

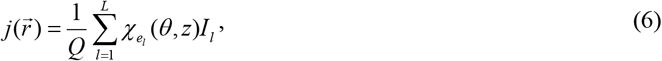

where *L* is the total number of electrodes; *e_l_* is the electrode *l*; *I*_l_ is the current applied to electrode *e_l_*; *Q* is the surface area of each electrode, assuming all electrodes have the same shape, and

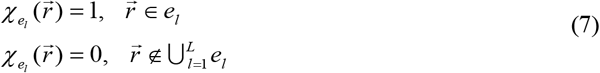

In this study, all of the electrodes have the same shape and dimension and are attached to the surface of the body. In the analytical solver, we assume the electrode has a rectangular shape with angular span of Δ*θ* and height of *E_h_*. The electrodes can all be placed in one layer or be split into multiple layers depending on the tradeoff between the area to be covered and the in-plane spatial resolution to be achieved. The area covered by the electrode *l* is

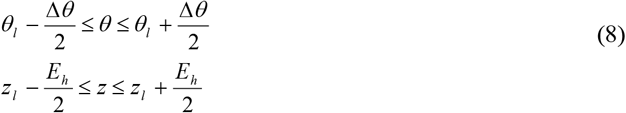

where (*r_0_*, *θ_l_*, *z_l_*) are the center of electrode *l*.

Let *E_a_* denote the area of an electrode. Substituting the current density (6) and potential (4) into the third boundary condition in (3), the Fourier coefficients *a_n_*, *b_n_*, *A_nm_*, and *B_nm_* can be computed as

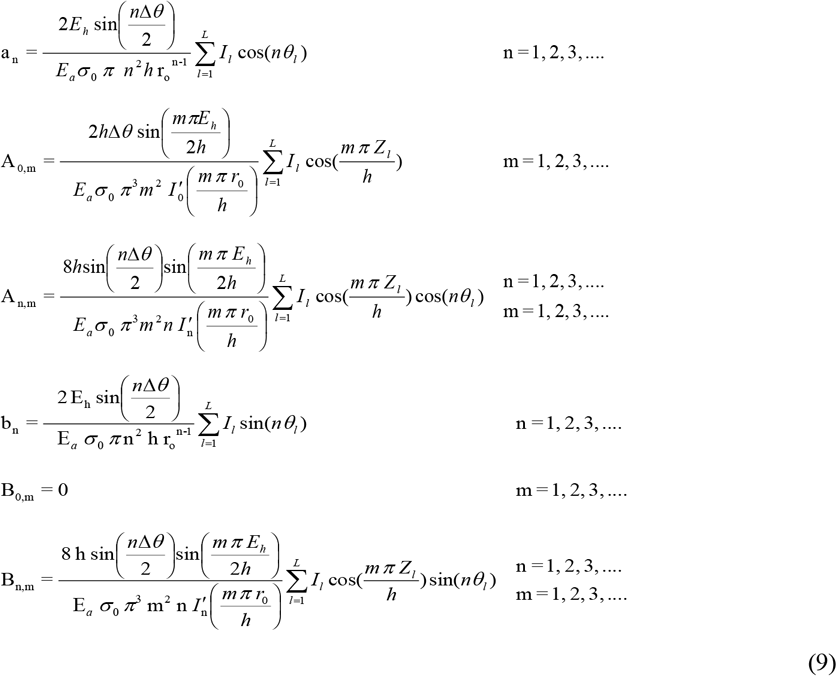

where *I*_n_′ is the first derivative of the modified Bessel function *I*_n_(*x*).

### 2.2 The 3D Reconstruction Algorithm

The reconstruction algorithm used in this study is based on a perturbation approximation which assumes that the conductivity distribution for which we are trying to solve differs only slightly from a uniform conductivity distribution by a perturbation *η*.

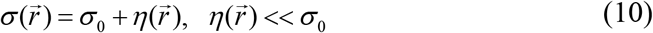

For a uniform conductivity, (1) becomes

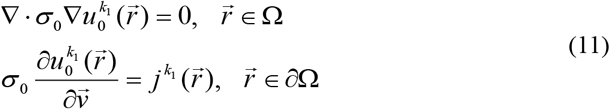

where *k*_1_ is the applied current pattern.

For the conductivity distribution *σ* in (10) and applied current pattern indexed by *k*_2_, linearizing the solution 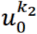 to (1) about *u*_0_ yields 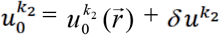 where 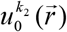 is the potential due to *σ_0_*, and 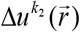 is O(*η)*. Then equation (1) becomes

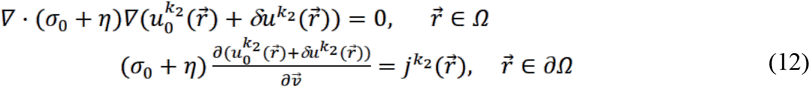

Combining (11) and (12), applying the divergence theorem, and neglecting the second and higher order terms, results in

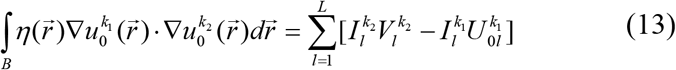

where 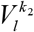 is the measured voltage on electrode *l* with conductivity distribution *σ* and current pattern *k*_2_, 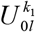 is the predicted voltage on electrode *l* with uniform conductivity *σ_0_* and current pattern *k*_1_. *K* is the total number of current patterns used. Denoting the area of an electrode by Q, where all electrodes are assumed to be uniform in size, using the ‘Ave-gap model’ [Cheng, 1989], we can make the approximations

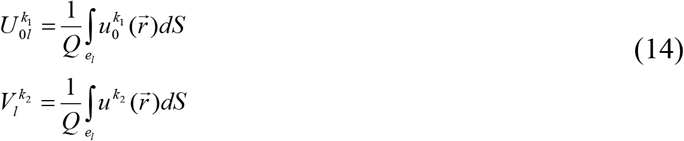

Discretizing the volume, (13) can be written as

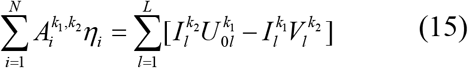

where *i* is the voxel index of the discretized volume, *N* is the total number of voxels, η_i_ is the perturbation in the i-th voxel, and

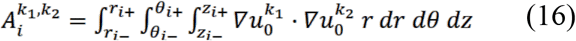

Thus, (15) can be written in a vector-matrix form

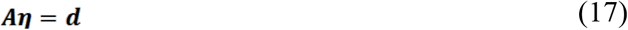

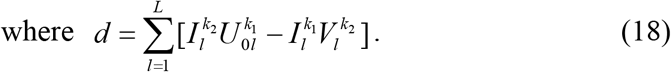

Here **d** is the data term, and the *K*_2_ by N matrix **A** is the Jacobian of the forward model with respect to a small perturbation in conductivity. The data term ****d**** can be calculated from the known current pattern, the measured voltage and the predicted voltage with a uniform conductivity, which is known or can be estimated as described in section 3. In time difference imaging, voltage taken at reference time *t_0_* will be used as 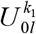.

As shown in (17), if **A** were non-singular, one could simply invert it and multiply by the data term to obtain *η*. Unfortunately, **A** is ill conditioned, and so it is necessary to regularize it to improve its conditioning in order to obtain a stable solution. The regularization of **A** will be described in section III. In this section the focus is to describe in detail how to compute the matrix **A** analytically.

The gradient of *u* can be computed from (4) and (9) as

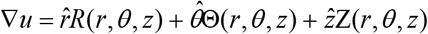

where

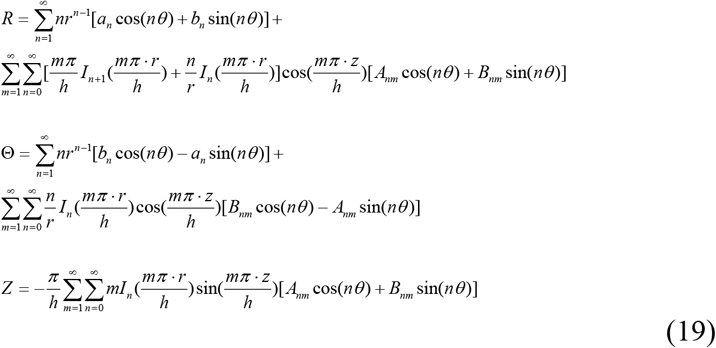

Adopting the notation

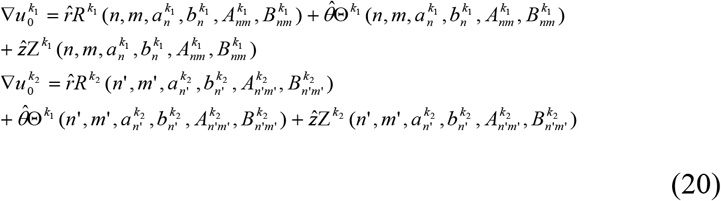

each entry of the matrix **A** is given by

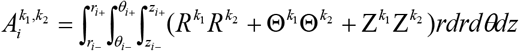

Solving the integrals results in

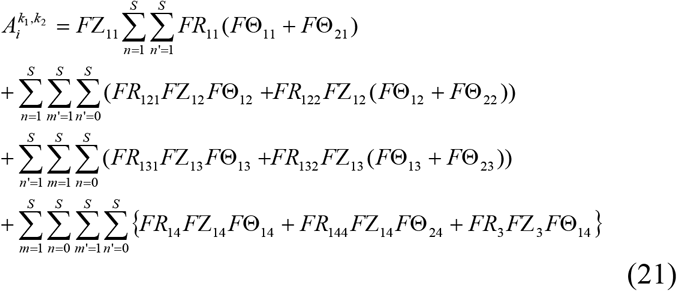

where

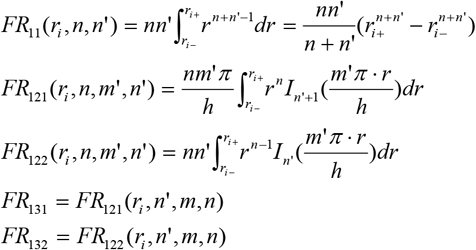

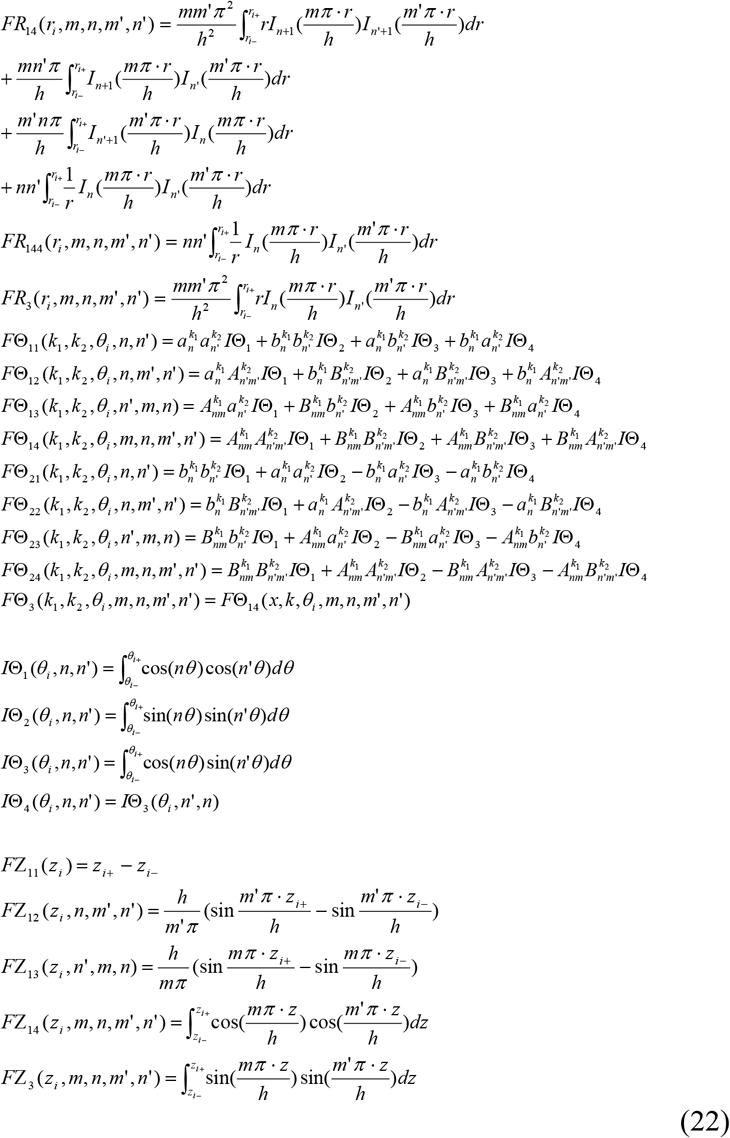

where *S* is the number of Fourier terms used in the approximation. The calculation of *FR*_121_, *FR*_122_, *FR*_131_, *FR*_132_, *FR*_14_ and *FR*_3_ must be done numerically using numerical quadrature. The others can all be solved analytically.

## 3 Reconstruction

The SMS-EIT GENESIS system is a real-time, 32-channel, 3D EIT system capable of running up to 20 frames per second at 10 kHz [Ashe 2014]. 31 trigonometric current patterns are applied simultaneously to all of the channels and the resulting voltages are measured on the electrodes. In the following experimental study, 32 electrodes were placed in 4 circular equally spaced rings. The current patterns and electrode configuration are presented in Figure 1 and Figure 2.

**Figure 1:**
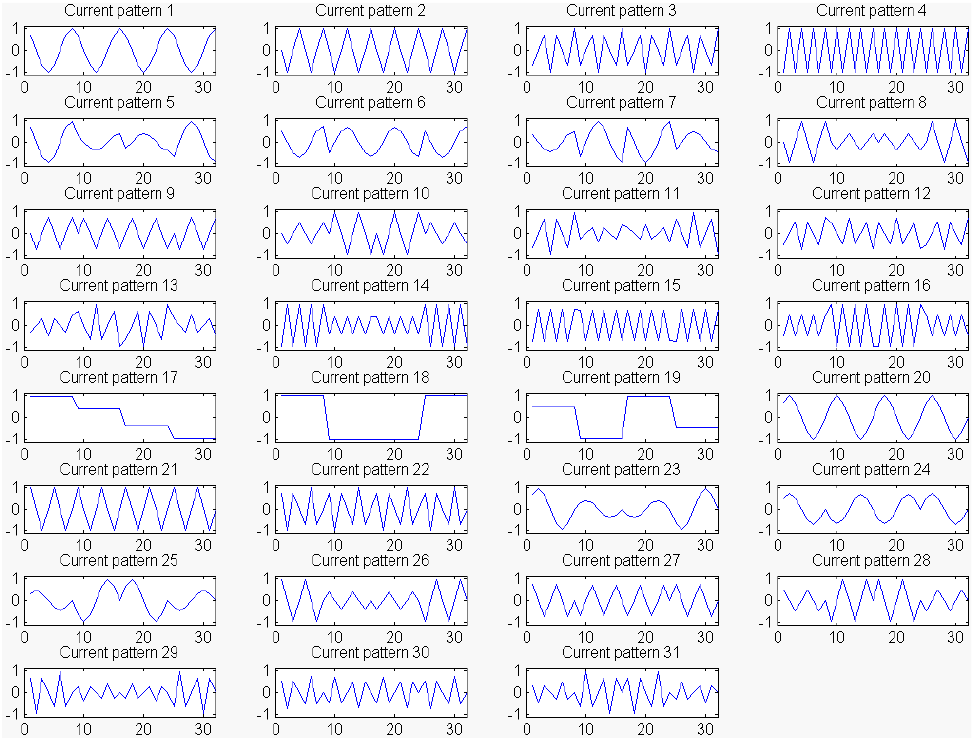
The 31 Applied Trigonometric Current Patterns.

**Figure 2:**
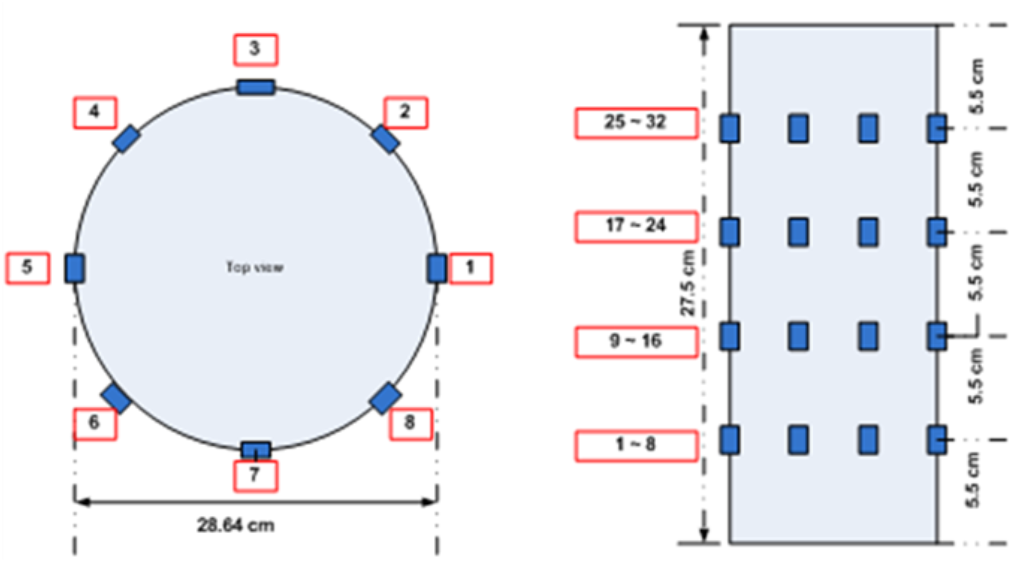
Electrode configurations.

A time difference imaging method is implemented in real-time on the prototype system. The voltage signal collected at a reference time *t*_0_ is used as a reference, which represents 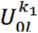 in (18) to calculate the data term *d*. The reconstructed image is the change in conductivity and permittivity compared to the reference time *t*_0_. The time difference imaging approach serves to reduce error caused by mismatch between geometry, electrode shape, location, and contact impedance in the model.

Note that a best fit constant-conductivity approximation to the measured data can be computed from

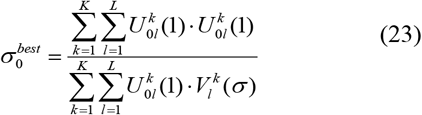

where 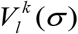 is the measured voltage signal at *t*_0_ and 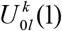 is the predicted voltage with a uniform conductivity of 1. One could then approximate the value of the conductivity by

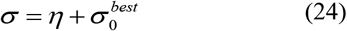

For a system with L electrodes and L-1 current patterns, there are L(L-1)/2 independent measurements, since the current-to-voltage operator is self-adjoint. In this work, the reconstructions are computed on the Joshua tree mesh [Cheney 1990], as shown in Figure 3, with 496 voxels, matching the number of degrees of freedom for 32 electrodes and 31 current patterns. Since the 32 electrodes were placed in 4 equally spaced rings, we used a 4-layer Joshua tree mesh with 124 voxels in each layer. The height of each layer is determined by the ring-to-ring distance between electrodes and the distance from the bottom / top of the cylinder to its closest ring of electrodes. For 2 rings of electrodes with 16 electrodes in each ring, a 2-layer Joshua tree mesh with 248 voxels in each layer is used.

**Figure 3.**
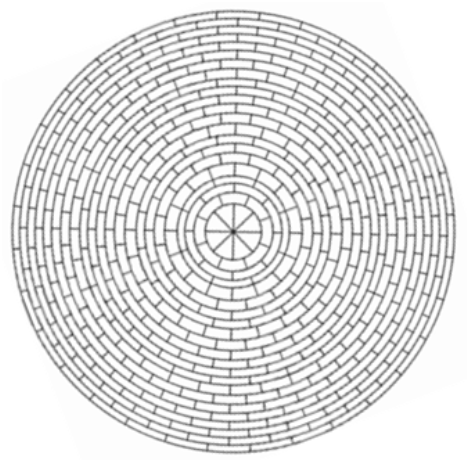
2D Joshua tree mesh with 496 voxels

The coefficient matrix **A** is pre-computed and saved to facilitate the real-time reconstruction. The matrix **A** is regularized using the same approach as in the NOSER algorithm [Cheney 1990]. That is, we solve the regularized system

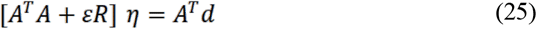

where *ε* is the regularization parameter, and *R* is a diagonal matrix with the diagonal terms of *A^T^A*.

The implementation of the reconstruction algorithm can be summarized as follows.

1. Pre-compute matrix **A** by equation (21);
2. Apply currents (total number of current patterns *K* = 31), and measure the resulting voltage on electrodes (total number of electrodes *L* = 32) at reference time *t*0 and at time *t*;
3. For each set of measured voltages, subtract a constant to ensure the voltages over all electrodes sum to 0;
4. Compute the data term ***d*** using (18), taking the measured voltage at reference time *t*_0_ as 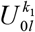;
5. Compute *η* using (25);

## 4 Validation of the Reconstruction Algorithm

### 4.1 Validation with Simulated Data

The forward solver from the open-source software package EIDORS [23] was used to generate data simulating a cylinder containing saline with conductivity 150 mS/m for the reference data and the cylinder filled with saline plus a spherical target placed near the third ring of electrodes from the top at the 7-8 o’clock position. The conductivity ratio between the spherical target and saline is 10:1. See Figure 4 for the target location and cylinder geometry.

**Figure 4:**
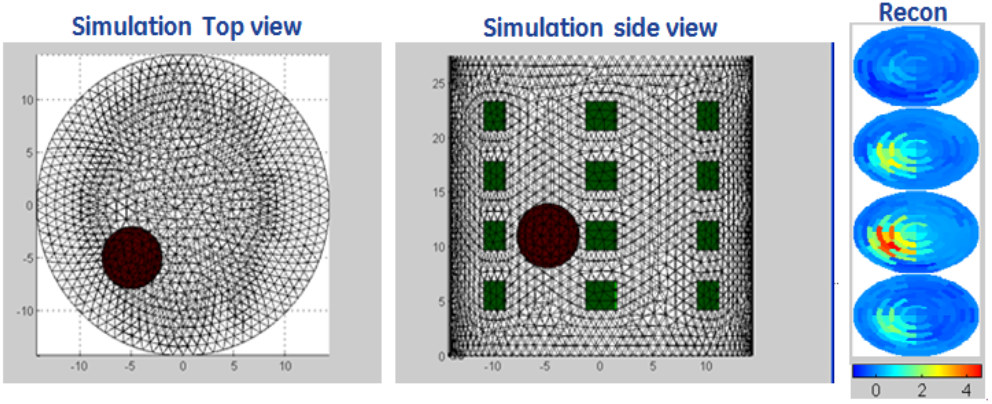
Simulation model used in EIDORS and the reconstruction image (Left: simulation top view, Middle: simulation side view, Right: 3D reconstructed images presented in 2D slides.)

At right in Figure 4 is the reconstructed time difference image at each layer in the mesh. Conductivity values higher than that of the reference image are displayed in red and lower values are displayed in blue. Conductivity units are arbitrary but consistent for a given electrode configuration and reconstruction. The image indicates a higher conductivity target at the 7-8 o’clock position in the 3rd layer from the top, which is consistent with the ground truth in both relative conductivity and position.

### 4.2 Validation with in-vitro phantom data

To validate the reconstruction algorithm with *in-vitro* phantom data, two sets of data were collected with the SMS-EIT system. Reference voltage data (a homogenous dataset) was collected on a cylinder tank filled with saline (150 mS/m). The inhomogenous dataset was collected after placing a small cylinder target in the saline tank, between the third and fourth ring of electrodes from the top, at the 2 o’clock position as shown in the left and middle figure of Figure 5. The 3D reconstructed conductivity image is displayed in 2D slices with 4-layer Joshua tree mesh which representing the electrodes layers. The reconstruction image indicates a higher conductivity target in the 2 o’clock position between the 3^rd^ and 4^th^ layer from the top.

**Figure 5:**
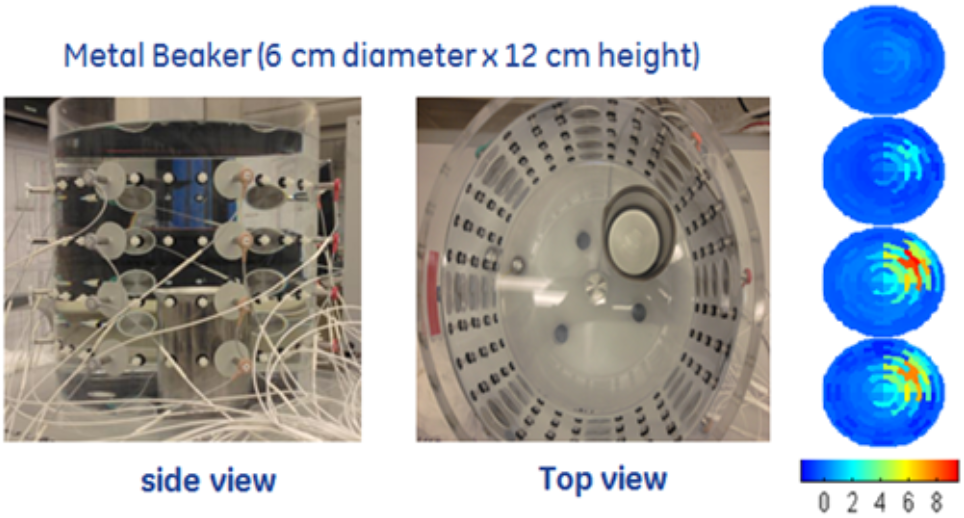
Phantom validation with a saline tank and a cylinder stainless steel target. (Left: side view, Middle: Top view, Right: 3D reconstructed images presented in 2D slides.)

### 4.3 Validation with Human Subject Data

The SMS-EIT system was validated under IRB approval to collect data and analyze system performance for healthy human subjects at GE Research. The system passed safety test to meet for IEC60601-1 safety requirement for medical device.

Commercially available ECG electrodes (Intelesens, Belfast, Ireland) with 3 cm conductive diameter were arranged in 4 layers and attached to the male subject’s torso as shown in Figure 6. The center-to-center distance between two neighboring electrode layers is 6 cm. The top ring was placed at the 3rd intercostal space and the bottom ring was placed at the bottom of the rib cage. A safety reference ground electrode was placed on the subject’s shoulder and connected to the system’s safety circuit to detect any unexpected leakage current. The system will switch off automatically if the leakage current is greater than the threshold of 0.450 mA_RMS_. A pulse oximeter, based on the photoplethysmograph, was placed on the subject’s index finger on the left hand, and data was collected while the subject breathed through a mask connected to a pneumotachograph to verify the impedance result of respiratory activity. The real-time 3D reconstructed image is shown in Figure 7, and the update speed is about 20 frame/second when displayed in 2D slices format.

**Figure 6:**
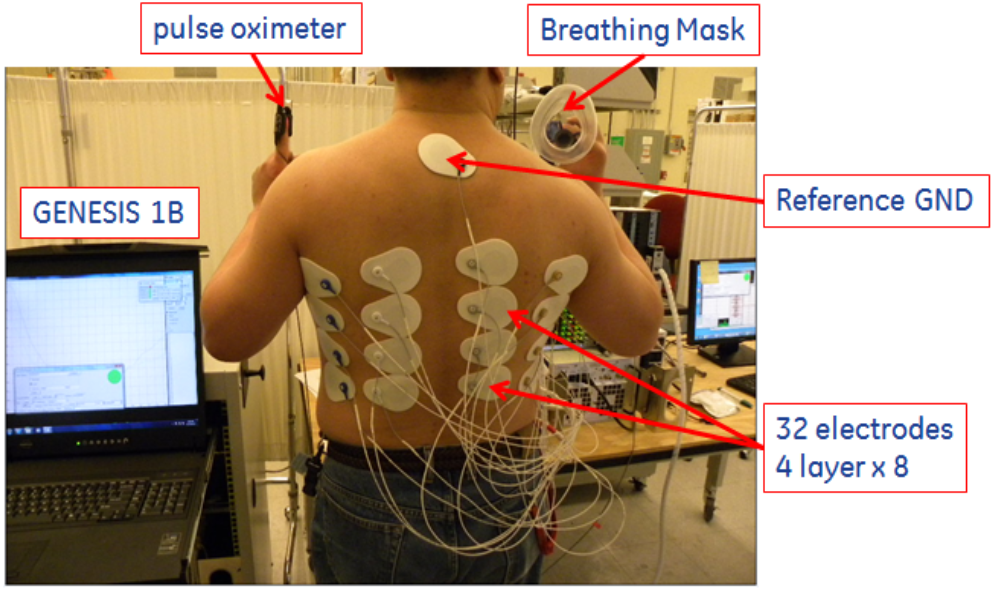
Experimental setup for human subject study.

**Figure 7:**
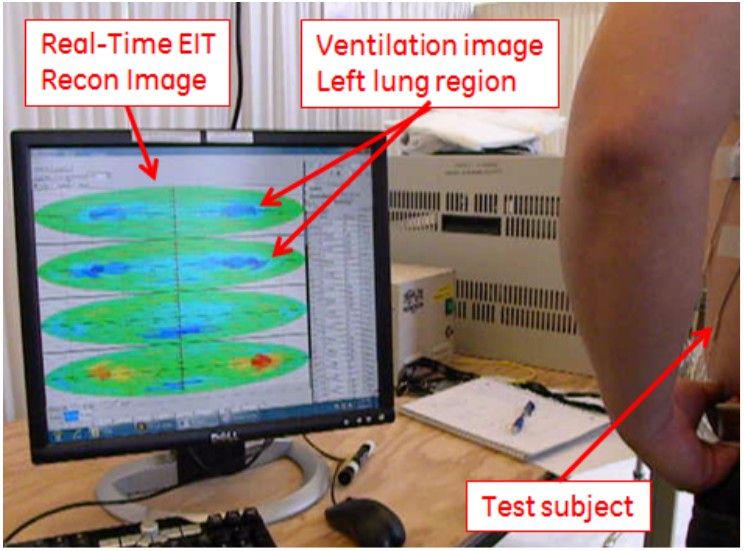
The real-time reconstructed image in a volunteer healthy subject.

The subject performed 1 minute of normal breathing, 1 minute of deep breathing, a 30 second breath hold, and 30 seconds of normal breathing in one 3-minute sequence, as shown in Figure 8. The waveform comparison between global conductivity, pneumotachograph, and photoplethysmograph has been previously reported in [Kao 2014].

**Figure 8:**
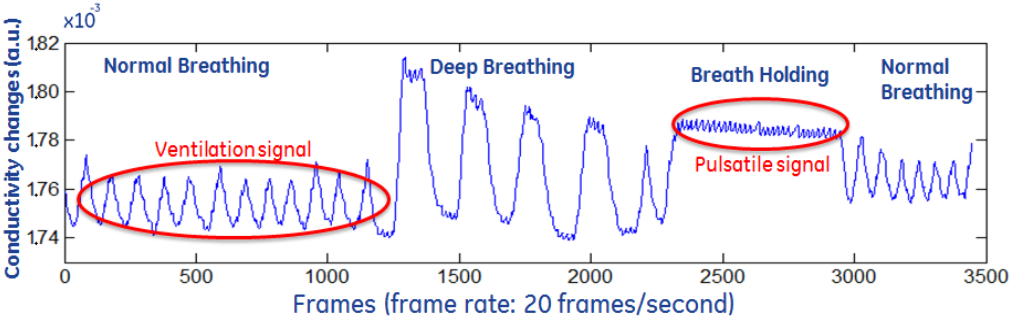
Detected ventilation signal and pulsatile cardiac activity signal on a healthy human subject study.

## 5 Lung Volume Estimates on Female Subjects

### 5.1 Study Design

This study was conducted in accordance with the amended Declaration of Helsinki. Data were collected on 6 female subjects in the EIT lab at Colorado State University, Fort Collins, CO under the approval of the IRB. Informed written parental consent and informed assent were obtained from subjects under age 18, and informed written consent from subjects age 18 and up were obtained prior to participation. Subjects ranged in age from 17 to 49 years, chest perimeters at the intra-mammary fold ranged from 29 to 34 inches, and bra cup sizes ranged from A to D.

### 5.2 Protocol

A breathing maneuver using a 2500 ml volumetric incentive spirometer (VIS) (Air-life 001904A by CareFusion) was used to record inspiratory volumes using the following protocol:

1. Subject exhales fully and places mouth tightly on the VIS
2. Slow inhale to a repeatedly comfortable level, desirably between 750 and 1000 ml
3. Breath-hold for about 3 s while the blue measurement guide on the VIS drops to the bottom
4. Inhale another additional 750 – 1000 ml
5. Breath-hold 3s for about 3 s while the blue measurement guide on the VIS drops to the bottom
6. Slowly exhale

Subjects were instructed in the procedure and asked to practice several times until they felt comfortable with the procedure. Two rows of 16 Philips 13951C pediatric ECG electrodes were applied around the circumference of the subject’s chest. Electrodes 1 - 16 were placed at the level of the intra-mammary fold, or vertebra T9, and electrodes 17 - 32 were placed above the breasts, at the level of vertebra T5. The circumference of the subject was measured at each row, and the average was used for the circumference of the cylindrical model. Biopac was used for 3-lead EKG measurements during data collection. 3D reconstructions were computed using the algorithm described in Section II.B.

### 5.3 Computation of lung volume estimates

Key time points in the maneuver were identified using a principal component analysis (PCA) of the data. From the PCA, a reference image was chosen representing a point of maximal exhalation prior to the start of the first inhalation maneuver. Frames corresponding to the end of the first inhalation, start of the second inhalation, and end of the second inhalation were also identified. These frames are denoted by

SFI = start of first inhalation
EFI = end of first inhalation
SSI = start of second inhalation
ESI = end of second inhalation.

Since the lungs become more resistive as they fill with air, a voxel was chosen to be in the lung region if it became more resistive from frame SFI to frame EFI or from frame SSI to ESI. Since the subjects in this study had healthy lungs, this criterion is not likely to incorrectly omit lung voxels but may include voxels corresponding to artifacts. From observations of the images, the outer 4 rows of voxels in the lower ring and the outer 6 voxels in the upper ring were omitted from the lung region. For most subjects this segmentation resulted in nearly all voxels in the upper layer except for the outer six voxel rings meeting the criterion to be included in the lung region. However, anatomically, this is reasonable since in most females, a cross-section at the T5 vertebrate level is above the heart but may include the aorta.

Ventilation volumes were computed using measures derived in [Muller 2015]. For each voxel *p* in the segmented lung region, an approximation to the local volume fraction of air, *f_a_*(*p*,*t*), is computed at time *t*. Denoting the reconstructed conductivity in voxel *p* at time *t* by σ_*a*_(*p*,*t*), and assuming that the conductivity changes in each voxel in the ventilation image sequence are caused by changes in volume of air in that voxel, the conductivity, σ_*a*_(*p*,*t*), is decomposed as

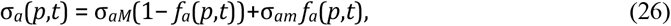

where σ_*aM*_ and σ_*am*_ are the maximum and minimum values, respectively, of the conductivity over the data collection period. From this decomposition, the volume fractions of air can be computed from the conductivities by

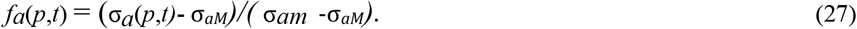

Denoting the volume of *p* by *vol*(*p*), the volume of air in voxel *p* at time *t* is estimated by

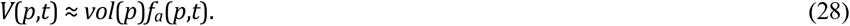

The cross-sectional area of a voxel was computed from the area of a pixel in the Joshua tree mesh, and the height of the voxel was taken to be one half the distance between the upper and lower rows of electrodes. Total lung volumes were then computed by summing over all voxels in the mesh.

### 5.4 Results

Table 1 contains the inhalation volumes recorded with the VIS. In the subsequent images, we show the results from Subjects 1 and 2 for brevity. The total lung volumes for each frame in the reconstruction computed from the segmented EIT images are plotted in Figures 9 and 10 for subjects 1 and 8, respectively with the key time points SFI, EFI, SSI, and ESI annotated in the figures. Plots for the remaining subjects are found in Figure 11.

**Table 1:** Inhalation volumes recorded using the volumetric incentive spirometer (VIS). Note that the subject held her breath between each inhalation while the VIS returned to 0 ml. Thus, the second inhalation represents volume inhaled in addition to the initial inhalation.

| Volume (ml) |  |  |  |
| --- | --- | --- | --- |
| Subject | Inhalation 1 | Inhalation 2 | Total |
| **1** | 750 | 1000 | 1750 |
| **2** | 625 | 625 | 1250 |
| **3** | 1000 | 1000 | 2000 |
| **4** | 250 | 500 | 750 |
| **5** | 1250 | 1250 | 2500 |
| **6** | 800 | 800 | 1600 |

**Figure 9:**
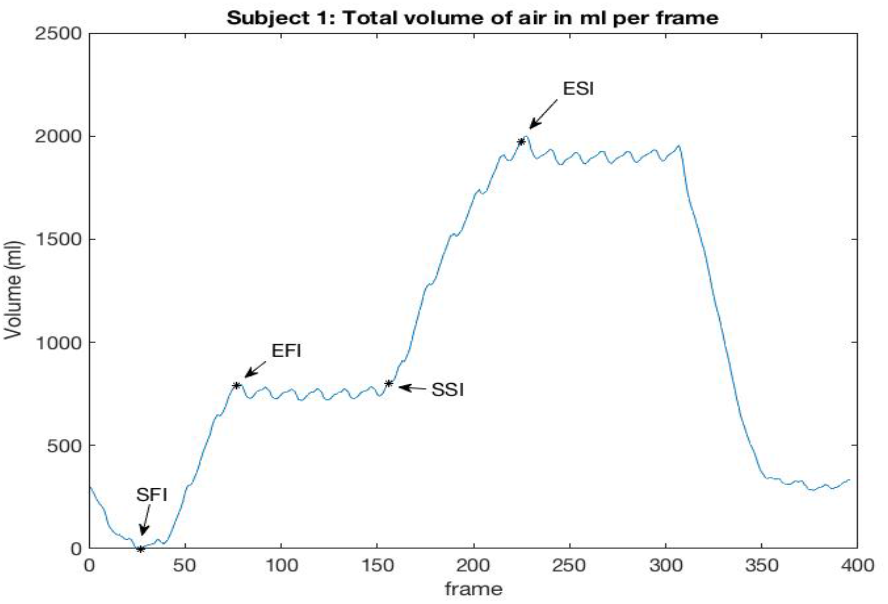
Lung volume per frame for Subject 1 computed from the segmented EIT images with key time points annotated.

**Figure 10:**
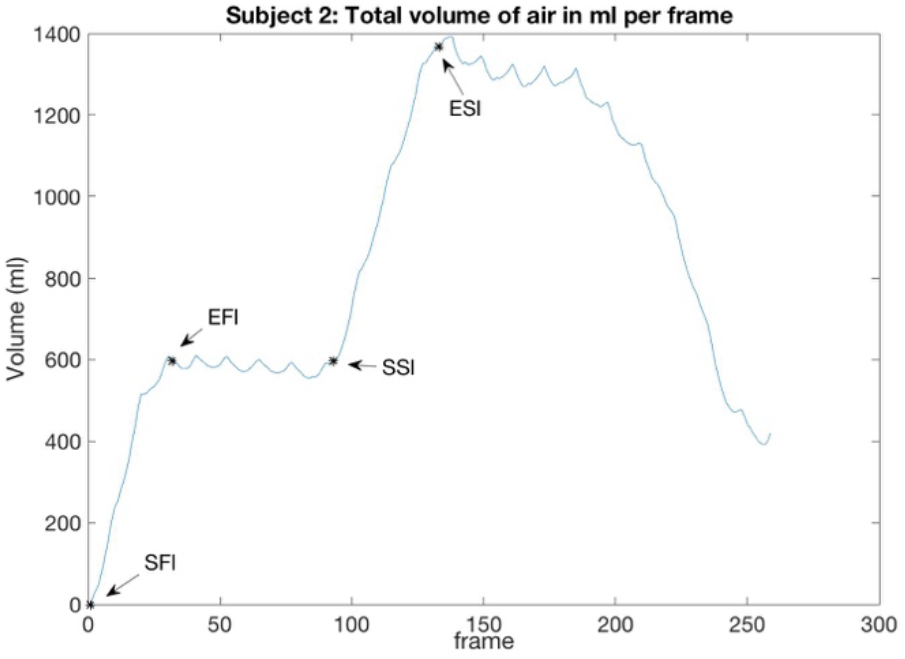
Lung volume per frame for Subject 2 computed from the segmented EIT images with key time points annotated.

**Figure 11:**
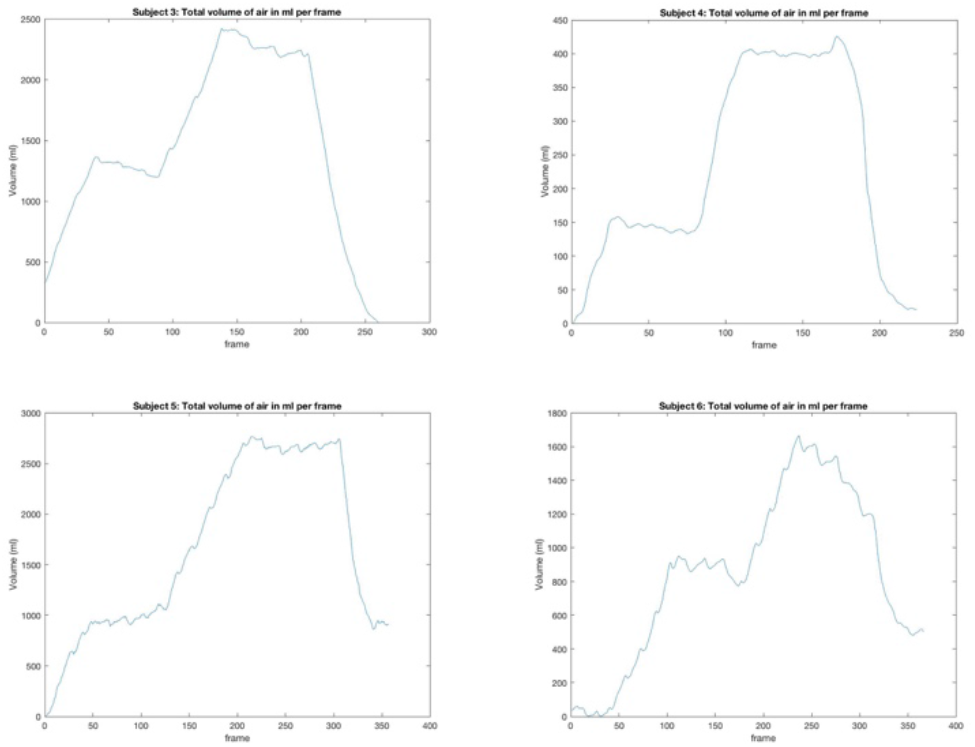
Lung volume per frame for subjects 3, 4, 5, and 6.

Figures 12 and 13 show the conductivity change in the segmented lung region from frame SFI to EFI (left), frame SSI to ESI (center), the overall change from frame SFI to ESI (right). Each image is shown on its own scale so that the changes are evident since the overall change has the highest dynamic range. All plots are in DICOM orientation. In Figure 12 one sees that the upper lung filled more than the lower lung during the first inhalation, and the lower lung was filled more significantly during the second inhalation, while in Figure 13, the lower lung fills more than the upper lung. This demonstrates the resolution of the algorithm along the cranio-caudal axis in addition to the in-plane resolution that is evident in the images.

**Figure 12:**
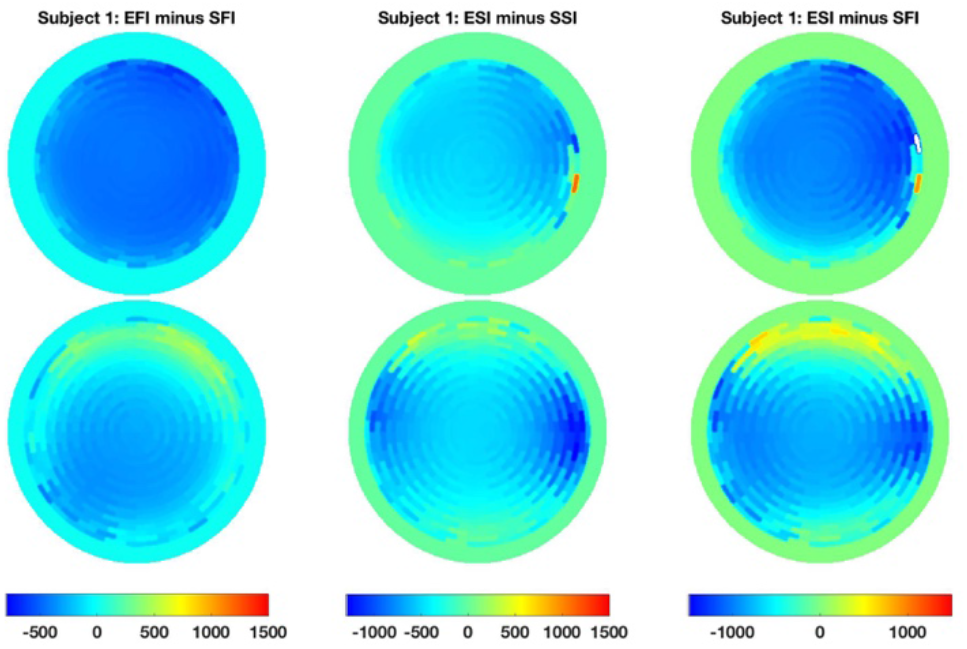
Ventilation difference images for Subject 1. Left: EFI minus SFI, Center: ESI minus SSI, Right: ESI minus SFI. The lower row of images depicts conductivity changes in the voxels centered on the intra-mammary fold, while the upper row depicts changes in the voxels centered on the ring of electrodes placed above the breasts.

**Figure 13:**
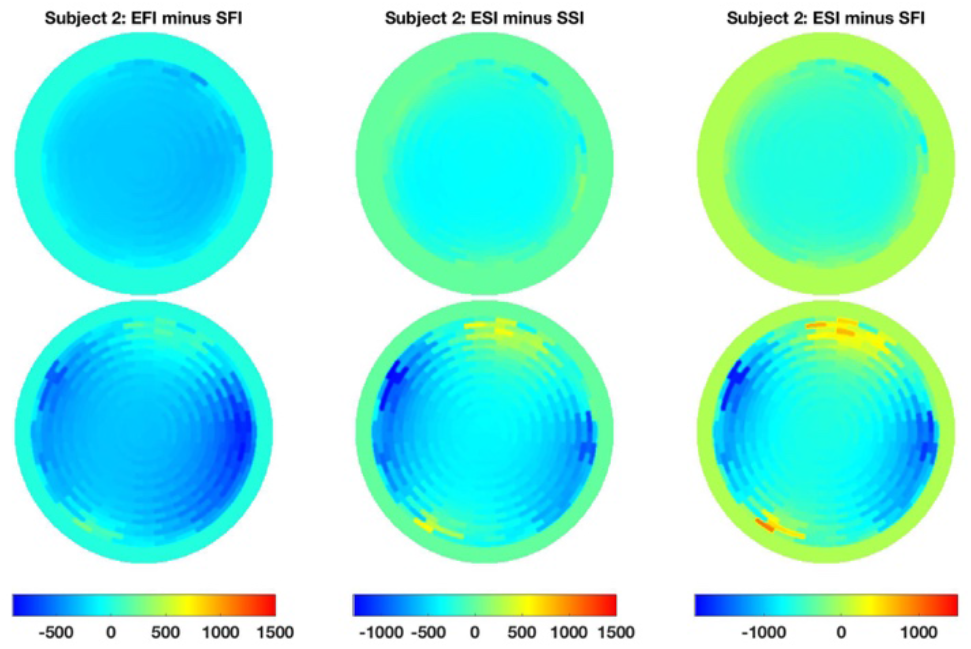
Ventilation difference images for Subject 2. Left: EFI minus SFI, Center: ESI minus SSI, Right: ESI minus SFI. The lower row of images depicts conductivity changes in the voxels centered on the intra-mammary fold, while the upper row depicts changes in the voxels centered on the ring of electrodes placed above the breasts.

Reconstructions of pulsatile pulmonary perfusion were computed for the periods of time during which the subject held her breath. Reconstructions from frames 190 to 243 for Subject 1 and frames 34 to 81 for Subject 2 were computed with frames 189 and 33 as a reference, for Subjects 1 and 2, respectively. Figures 14 and 15 show a single frame in the sequence of reconstructed images that captures the systolic heart. In each figure the time trace of the conductivity heart voxel is shown in red, and time traces of the conductivity of a voxel from the left lung and right are shown in blue and green, respectively.

**Figure 14:**
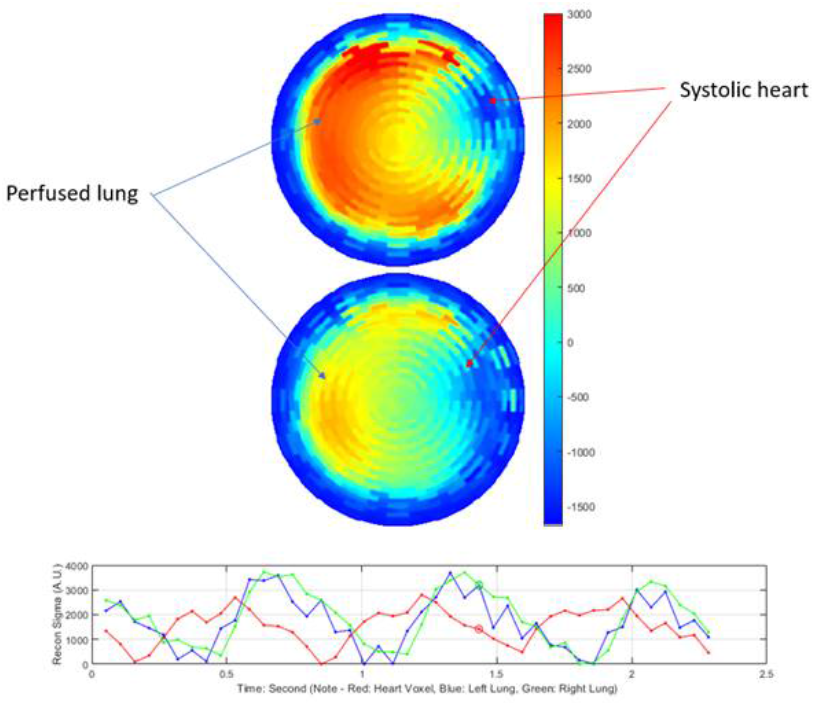
Time difference image for one frame in the pulsatile pulmonary perfusion sequence for Subject 1. The frame is indicated by a red star in the lower plot which depicts the time traces of the reconstructed conductivity in a voxel from the heart region (red line), left lung (blue line), and right lung (green line). Plots are in DICOM orientation.

**Figure 15:**
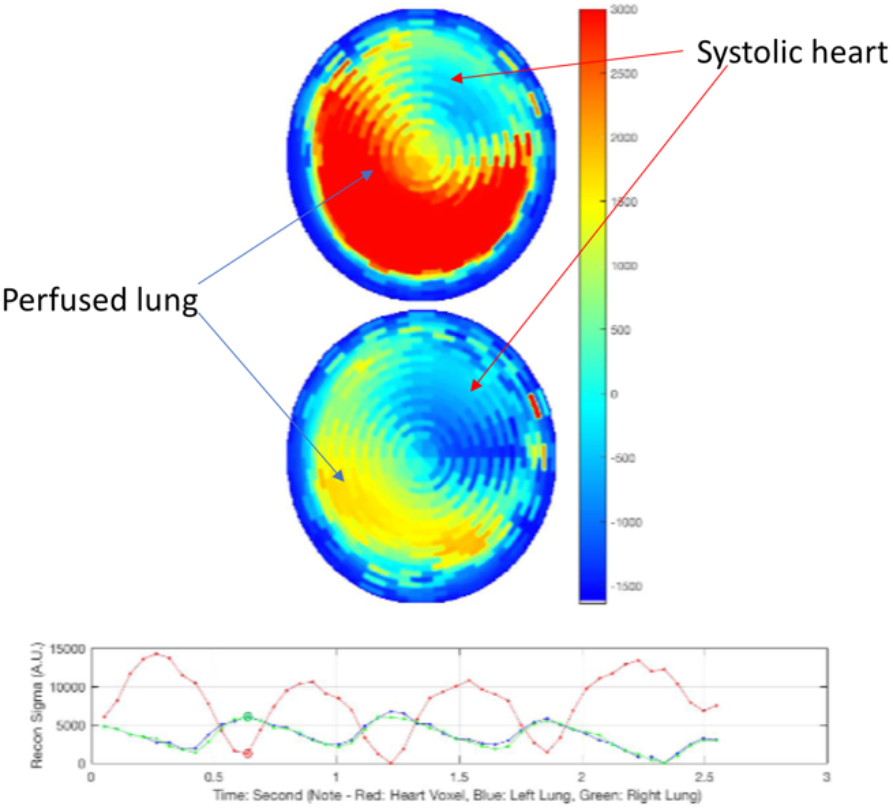
Time difference image for one frame in the pulsatile pulmonary perfusion sequence for Subject 2. The frame is indicated by a red star in the lower plot which depicts the time traces of the reconstructed conductivity in a voxel from the heart region (red line), left lung (blue line), and right lung (green line). Plots are in DICOM orientation.

## 6 Conclusion

This paper provides a detailed derivation of the analytical forward solution and the Jacobian matrix for a real-time implementation of a 3-D reconstruction algorithm based on the ToDLeR algorithm. The algorithm is validated with simulated data from EIDORS and *in-vitro* data collected from a saline tank. Two human subject studies are presented, demonstrating that the prototype SMS-EIT system has the ability to detect pulmonary ventilation and pulsatile cardiac activity signals in real-time.

## 6 Acknowledgment

We thank Dave Davenport, Jeff Ashe, Jim Sabatini, David Shoudy, Gregory Boverman and Xing Wang from GE Research who helped to build the EIT prototype system.

